# lncRNA-mRNA network analysis reveals novel biomarkers in prostate adenocarcinoma

**DOI:** 10.1101/2023.08.16.553489

**Authors:** Houri Razavi, Ali Katanforoush

**Affiliations:** Department of Data and Computer Sciences, Shahid Beheshti University, Tehran, IRAN

**Keywords:** Prostate adenocarcinoma, lncRNA, survival analysis, cancer hallmarks, network analysis

## Abstract

Prostate adenocarcinoma (PRAD) is the second most frequent cancer among men worldwide. lncRNAs have been suggested as novel promising biomarkers for cancer diagnosis and prognosis and therapeutic targets. Despite the previous investigations, still, comprehensive studies are required to discover the regulatory roles of lncRNAs in PRAD.

We introduced a PRAD lncRNA-mRNA network (PLMNET) to identify novel PRAD biomarkers and therapeutic targets among lncRNAs. The PLMNET is constructed based on RNAs that are differentially expressed and had strongly correlate with each other. PANTR1, HOXB-AS3, EMX2OS, GATA3-AS1, LINC01116, LINC02385 are PLMNET hubs that are considered key regulatory lncRNAs of PRAD. We also mapped PLMNET lncRNAs to their associated GO-terms and hallmarks. Literature review confirms our suggested biomarkers and their associated hallmarks.

The high degree nodes of PLMNET, GATA3-AS1, EMX2OS, LINC01116, LINC02137, RP11-431N15.2, LINC01671, LINC02806, ENSG00000231196, ENSG00000244252, RP11-379F12.3, and SLC2A9-AS1 indicate significant association with overall survival. In addition, we found that the hubs’ regulatory roles have previously been confirmed in different types of cancers by literature. For example, PANTER hub exhibited a high expression correlation with three known tumor suppressors (SKI, SKP2, and SH2B3) and is associated with “Sustaining proliferative signal”. The literature has confirmed that PANTER may promote cancer cell proliferation in prostate carcinoma. It also promotes renal cell carcinoma and Hepatocellular carcinoma progression. Furthermore, literature review proves that EMX2OS and LINC01116 are key regulatory lncRNAs of prostate adenocarcinoma and HOXB-AS3 and GATA3-AS1 have regulatory roles in other cancer.

According to pathway analysis, PLMNET hubs seem to contribute to PRAD proliferation, invasion, metastasis, and apoptosis via dysregulation of anaphase promoting complex/cyclosome (APC/C) subunits and dysregulation of mitotic cell cycle.

These results introduce PLMCNET hubs as potential molecular biomarkers for diagnosis and prognosis of PRAD and suggest therapeutic targets for PRAD patients.

## 1. Background

Prostate cancer is the second most prevalent cancer in men and second leading cause of cancer death in men all over the world [1]. Its high mortality rate is associated with lack of specific and sensitive methods for early prostate cancer screening and lack of effective therapies [2]. Therefore, it is necessary to further explore the pathogenesis of prostate cancer and develop therapeutic targets for prostate cancer patients [3,4].

Differential expression analysis is a general approach to find disease-associated biomarkers detects biomarker by screening gene expression changes between normal and disease groups. A drawback is that the approach considers the individual changes within a single family of biomolecules like lncRNAs or mRNAs. In contrast, in differential network analysis, a network is built based on expression correlation then candidate biomarkers are chosen by investigating topological features of the network. Although correlation tends to generate complicated networks, biomarker candidate selection only based on network topology ignores the changes on a single biomolecule level [5]. It seems that biomolecules with strongly altered connectivity between distinct biological groups might play a vital role in the disease under study [6]. Differential expression and differential network analysis look into data from two separate but complementary perspectives: the former focuses on the change of a single biomolecule while the latter concerns the change in pairwise associations between biomolecule pairs[7]. Here, we apply a combination of both models to gain their advantages. We seek for the critical lncRNAs playing role in PRAD pathways through a network-based analysis. We hypothesized that hub RNAs could have a vital regulatory role in cancer progression and treatment and be cancer biomarkers or/and therapeutic targets.

lncRNAs are regulators of gene expression and play critical roles in shaping cellular activities. They can modulate chromatin function, regulate the function of membraneless nuclear bodies, change the stability and translation of cytoplasmic mRNAs and interfere with signaling pathways. Recent studies have shown that they mediate oncogenic or tumor-suppressing effects [8]. They affect gene regulatory mechanisms and play a critical role in cellular homeostasis, including proliferation, invasion, migration, metastasis, and apoptosis [9,10]. As a result, hundreds of researchers are focused on lncRNAs as novel biomarkers or therapies, yet we are at the beginning.

Recent studies analyzed lncRNA-mRNAs ceRNA networks via the expression correlation with miRNAs [11]. They refused to directly study the correlation between lncRNAs expression and mRNAs expressions for network construction. However, lncRNAs can adjust mRNA stability through different mechanisms: 1) Direct interplay with miRNA or RBP binding sites in target mRNA; 2) Suppression of miRNAs or RBPs to keep away from interplay with mRNAs; 3) Acting as scaffolds to facilitate RBP-mRNA interplays; 4) Interplay with m6A machinery to adjust m6A levels of target mRNAs [12]. In fact, mRNA modulation with miRNA intervention is a subset of lncRNAs regulatory roles. In order to consider all regulatory mechanisms of lncRNAs at once, we directly compute the expression correlation among lncRNAs and mRNAs.

Furthermore, lncRNAs have been involved in the acquisition of every hallmark of cancer cells, from the innate capacity of proliferation and survival, through increased metabolism, to the relationship with the tumor microenvironment. Evidence proved that lncRNAs transcriptionally regulate key oncogenic or tumor-suppressive transcription factors such as p53, MYC, the estrogen receptor or signaling cascades such as the Notch pathway. For example, *MEG3 lncRNA participates* in the p53 regulatory net without being transcriptional targets of p53. *MEG3* contains an evolutionary conserved RNA structure that mediates p53 activation in *trans and is* downregulated in multiple cancers. Numerous lncRNAs are either regulated by or regulate the expression of the proto-oncogene MYC[10]. These findings legitimize the possibility of associating lncRNAs to cancer hallmarks via mRNA expression correlation. Juan and her colleagues [13] introduced a model that maps miRNAs to mRNAs and cancer hallmarks. They suggest that RNA crosstalk is not only of fundamental importance in physiological conditions, but is also crucially relevant in various cancers. Their analyses and validations demonstrate how the cancer-associated RNA regulatory network can be used to accelerate the discovery of RNA-based biomarkers and potential therapeutics. In this research, we mapped lncRNAs to the relavant Gene Ontology (GO) terms and cancer hallmarks to better understand the regulatory roles of lncRNAs in PRAD.

First, a PLMNET was constructed based on differentially expressed RNAs among normal and tumor samples. Then the most expression correlated RNAs were selected to identify PRAD-related lncRNAs. lncRNAs mapped to related cancer hallmarks via connecting their related mRNAs to relevant GO terms. The results prove that hubs of the PRAD network were cancer-related lncRNAs and introduced as biomarkers or/and therapeutic targets in PRAD.

## 2. Material and methods

### 2.1. RNA expression profiles

The mRNA expression profiles of PRAD solid tissues were downloaded from Cancer Genome Atlas (TCGA) [14] using the TCGABiolinks package [15]. For mRNA expression data, gene expression was measured through mapping RNA-Seq by Expectation Maximization (RSEM). Initial data consists of 19947 mRNA expressions from 554 samples where 502 were tumor tissues and 52 were adjacent non-cancerous tissues. First, we removed genes with RSEM expression values of 0 in all samples and then log 2-transformed the expression levels. PRAD lncRNAs expression was directly obtained from the Atlas of Noncoding RNAs in Cancer (TANRIC) [16]. In total, 12 727 lncRNAs were quantified for the expression levels as reads per kilo base per million mapped reads (RPKM). Finally, 426 samples, 52 normal and 374 tumors, with both lncRNA and mRNA expression profiles were selected for analysis. lncRNAs were displayed by their ensemble id in TANRIC, so the Ensemble database was used to convert the ensemble ids to gene symbols. Some lncRNAs present by their ensemble id since they are novel transcripts and don’t have a gene symbol.

### 2.2. Differential expression

EdgeR package was used to identify the differentially expressed mRNAs and lncRNAs between the normal and tumor samples. Most significant RNAs with FDR < 0.05 and |logFC| > 2 (361 mRNA and 54 lncRNA) were selected as initial nodes of network.

### 2.3. PRAD lncRNA-mRNA network (PLMNET)

Initial PLMNET was constructed by the differentially expressed mRNAs and lncRNAs as initial nodes. To define PRAD edges, we computed the Pearson correlation coefficient (R) of each RNA pair. All RNA pairs with R> 0.9 and p-value < 0.05 were selected as final edges. For each node in PLMNET, node degree is defined as the number of edges incident to it.

Finally, nodes with zero degrees were omitted from the network. The final PLMNET includes 23 lncRNAs and 52 mRNAs.

### 2.4. Calculating topological network properties

The topological features of PLMNET, including degree and Betweenness centrality (BC) calculated by the “igraph” package in R. Node degree, is the number of edges incident to the node. The BC indicates the influence of a node over the flow of information in the network. High BC represents the key regulatory role of the node in the network. We also detected the cancer-related RNAs by searching for the research papers that discovered a relationship between that RNA and cancer in PubMed.

### 2.5. Pathway enrichment analysis

PLMNET mRNAs pathway enrichment analysis at the REACTOME and KEGG pathways was performed using DAVID Bioinformatics Resources (http://david.abcc.ncifcrf.gov/). The DAVID enrichment analysis was limited to KEGG pathways with the whole human genome as background. Functional categories with *p*-value of < 0.05 and an enrichment score of > 1.5 were considered statistically significant. Finally, Reactome results illustrate due to its complete and precise data on our desired input [17].

### 2.6. Network visualization

PLMNET were visualized by Cytoscape 3.8.2, and topology analysis was performed by the package of ‘igraph’ in R language.

### 2.7. Survival analysis

The samples were classified as “Low count” and “High count” based on their lncRNAs expression. Classification threshold defined by *surv_cutpoint* function of *survminer* package. Statistical significance was set at p-value < 0.05. Then Kaplan–Meier and log-rank test was used to determine the association between the overall survival (OS) and PLMNET lncRNAs.

### 2.8. Construction of PRAD hallmark associated RNA network (PHANET)

To investigate the function and effect of PLMNET lncRNAs in cancer, first, the mRNAs mapped to their GO terms of their biological process using DAVID Bioinformatics Resources [18]. Chen et al. [19] review was used for mapping the GO terms to their related cancer hallmarks. We also consider the expression correlation among lncRNA and mRNAs. Finally, PHANET depicts a flowchart of connections among lncRNAs, mRNAs, GO terms, and cancer hallmarks in PRAD.

### 2.9. Literature review

We used Cosmic cancer gene census and PubMed to detect oncogenes, tumor suppressor mRNAs of the PRAD network. We also applied PubMed search engine to review the related literature on each significant lncRNA. The complete workflow represents in Fig 1.

**Fig 1.**
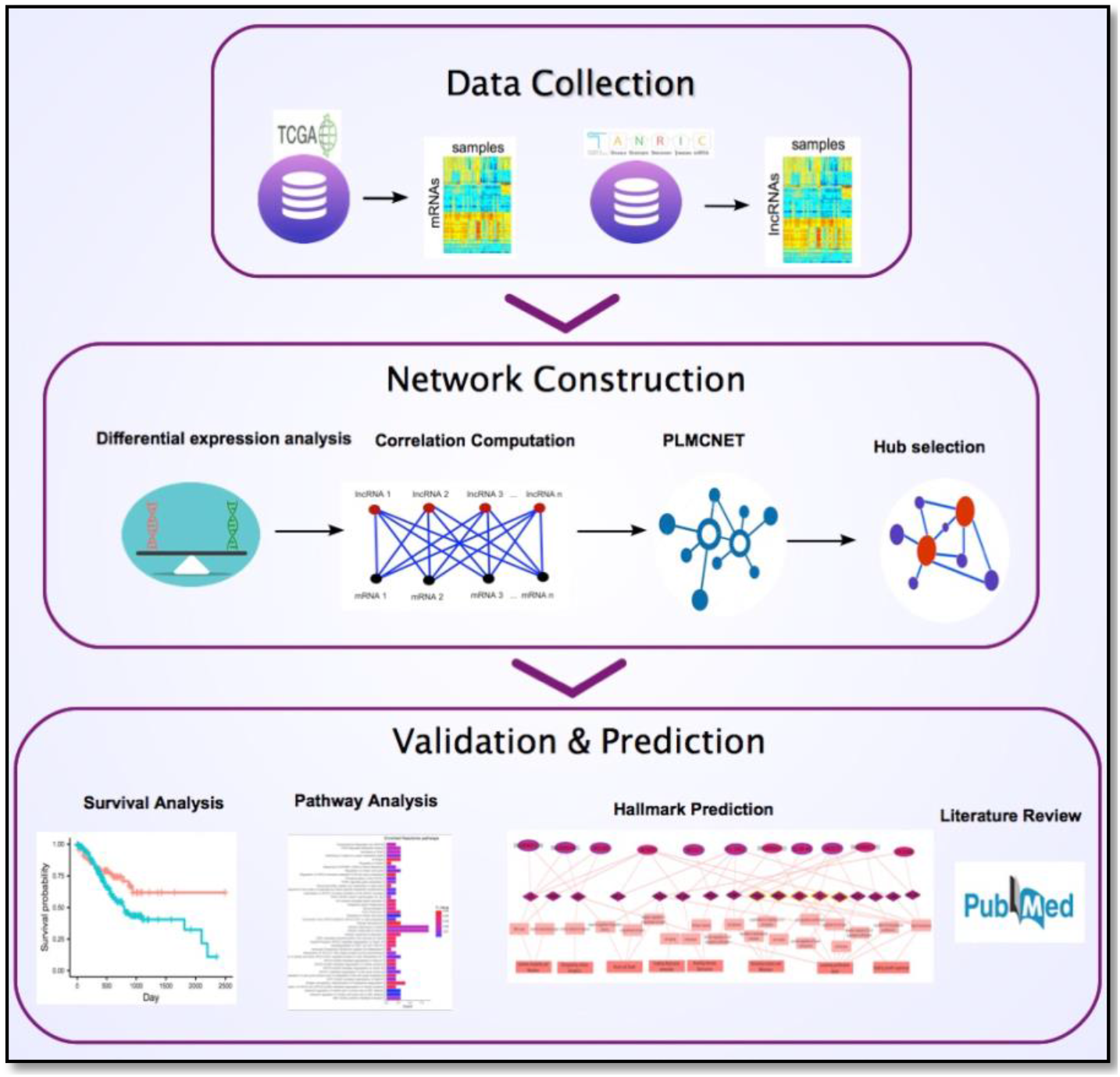
Workflow of PRAD key regulatory lncRNA detection

## 3. Results

### 3.1. Detecting differentially expressed mRNAs and lncRNAs

Differential expression analysis detected 8872 mRNAs and 1966 lncRNAs differentially expressed between PRAD normal and tumor samples. Then the most significant differentially expressed RNAs (with P < 0.05 and |logFC| > 2) were selected as initial nodes of PRAD lncRNA-mRNA network (Supplementary file1, 2). A total of 361 differential expressed mRNAs (191 up- and 170 down-regulated) and 57 lncRNAs (23 up-, and 34 down-regulated) were detected between PRAD and normal samples. Volcano plots displaying the distribution of PRAD lncRNAs and mRNAs expression provided using the R packages ggplot2 (Fig 2).

**Fig 2.**
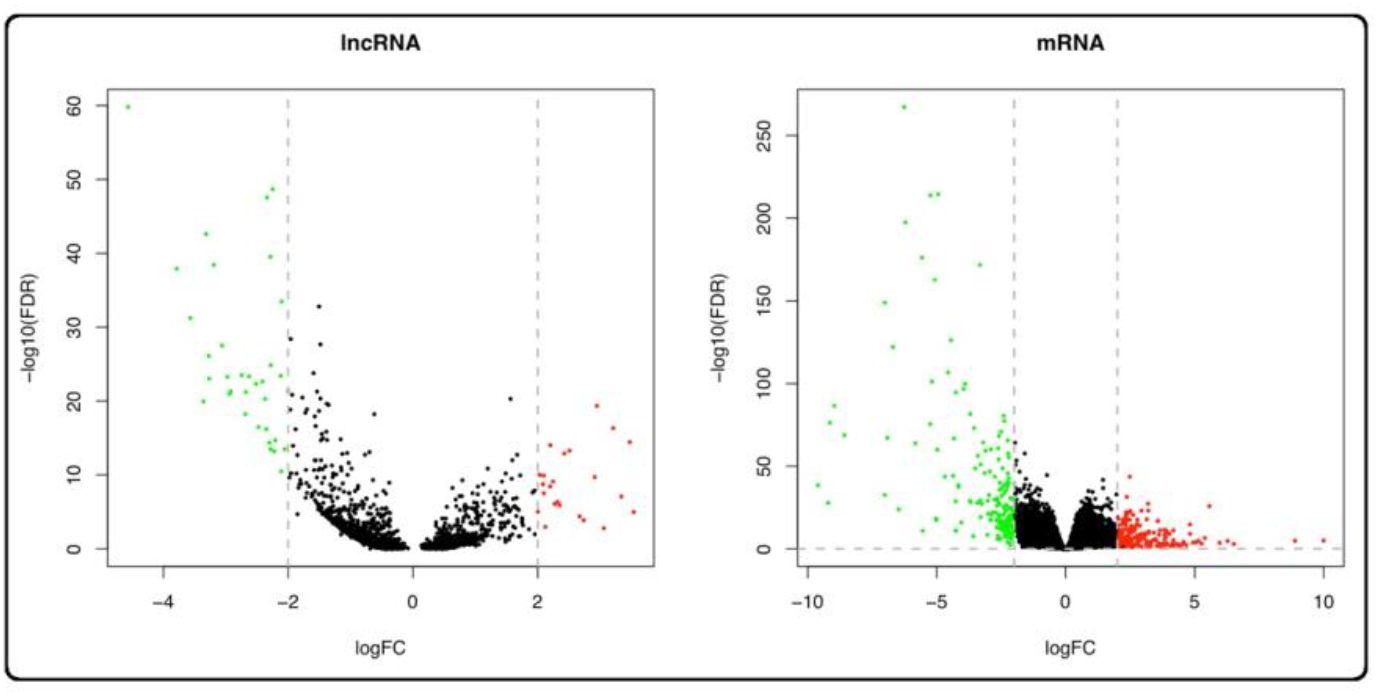
PRAD lncRNAs and mRNAs expression Volcano plots. X-axis indicates the expression differences of mRNAs, and lncRNAs between PRAD and normal samples, and Y-axis represents log transformed false discovery rate (FDR) values. Red dots represent the up-regulation, and green dots represent down-regulation.

### 3.2. PRAD lncRNA-mRNA network

lncRNAs and mRNAs with significant differential expression were considered as initial nodes of PLMNET. The network’s edges were defined based on a high Pearson correlation among RNAs expression. Nodes with zero degrees were omitted from the network since they hadn’t a strong correlation with others. The final network with 23 lncRNAs and 52 mRNAs depicts in Fig 3 (DDHD2 and SCARNA10 omit in Fig 3 since they made a separate component with only one edge). As seen in Fig 3, The high degree nodes of PLMNET are lncRNAs. For example, PANTR1 lncRNA with the highest degree among all nodes of PLMNET had a strong correlation with 32 mRNAs which all had significant differential expression among normal and tumor samples. A list of PLMNET edges with their correlation and p-values is in Supplementary file3.

**Fig 3.**
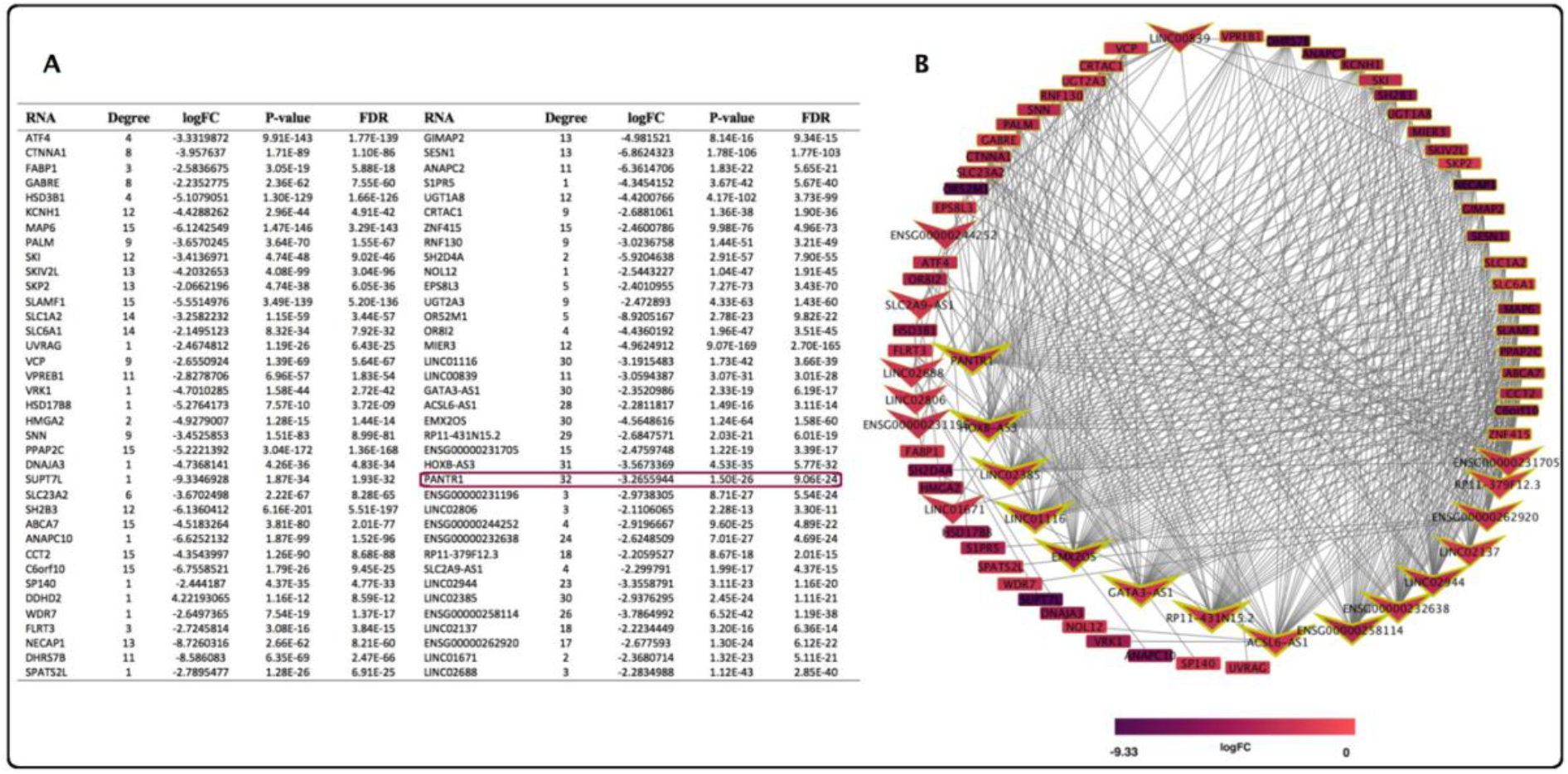
PRAD mRNA-lncRNA network. (**A**) network nodes and their characteristics (**B**) mRNAs represent by rectangles, and lncRNAs represent by arrows. Node filling color indicates log fold change of differential analysis among tumor and normal samples. The node’s border thickness represents the node degree. Nodes’ degree decrease by counterclockwise rotation.

### 3.3. Topological features of PLMNET

PLMNET includes 75 nodes and 411 edges. Among the 75 nodes, 52 (69.3%) are mRNAs and 23 (30.7%) are lncRNAs. 94% of PLMNET mRNAs and 30 % of lncRNAs are cancer-related while 36% of these lncRNAs don’t have a name since they are novel RNAs or their exact function is not clear. 13% of these mRNAs are oncogenes or tumor suppressors (CTNNA1, SKI, SKP2, HMGA2, SH2B3, SPATS2L, DDHD2, Fig 3b). The lncRNAs nodes have higher degrees, betweenness, and logFC than mRNAs nodes in the PLMNET (avg. 18.68 vs. 7.9 for degrees, Figure 4a: A; avg. 120.68 vs. 30.8 for betweenness, Figure 4a: B; avg. −2.83 vs. −4.23 for logFC, Figure 4a: C). According to network analysis principles, we expect that nodes with the highest degree and betweenness have a key regulatory role in PRAD initiation or progression. Testing this hypothesis, PLMNET nodes were sorted in descending order according to their degrees. We chose the top 8 percent of RNAs with the highest degrees as the hub components. PANTR1, HOXB-AS3, EMX2OS, GATA3-AS1, LINC01116, LINC02385, the hubs of PLMNET, are all lncRNA. After PubMed investigation, we got that all these hubs except LINC02385 are cancer-related [20].

**Fig 4.**
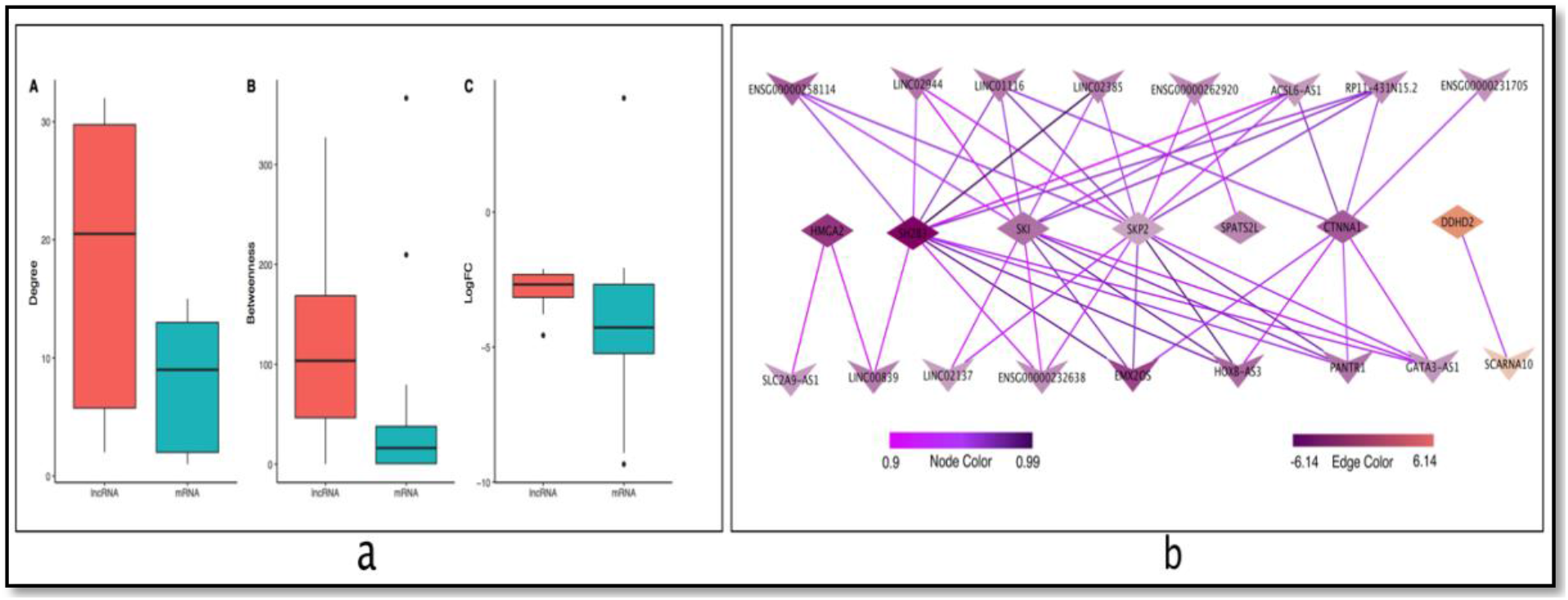
**a.** The distinct degree, BC, and logFC between lncRNAs and mRNAs in PLMNET. **b**. Correlation among PLMNET oncogene and tumor suppressor mRNAs and lncRNAs. Nodes color represents logFC and edges color represents their expression correlation.

Furthermore, VCP has the highest BC among PLMNET RNAs which implies its key regulatory role in PRAD. VCP is related to DNA repair process and promotes instability and mutation [21]. It has a strong expression correlation with 5(83%) hubs of PLMNET. PLMNET nodes list with degrees, BCs, logFCs, and p-values present in Supplementary file4.

### 3.4. PLMNET pathway analysis

A total of 41 Reactome pathways were significantly enriched (p-value <0.05) for PLMNET mRNAs (Supplementary file5), depicted in Fig 5 [22]. “*Aberrant regulation of mitotic exit in cancer due to RB1 defects”* was the most significant pathway of PLMNET. The enriched pathways are related to regulation of mitotic process, dysregulation of Anaphase promoting complex (APC) substrates, regulation of mitotic cell cycle, transcriptional regulation, Cellular responses to stimuli, Synthesis of DNA, and TP53 Metabolic Genes regulation. All the enriched pathways are well known to contribute to carcinogenesis.

**Fig 5.**
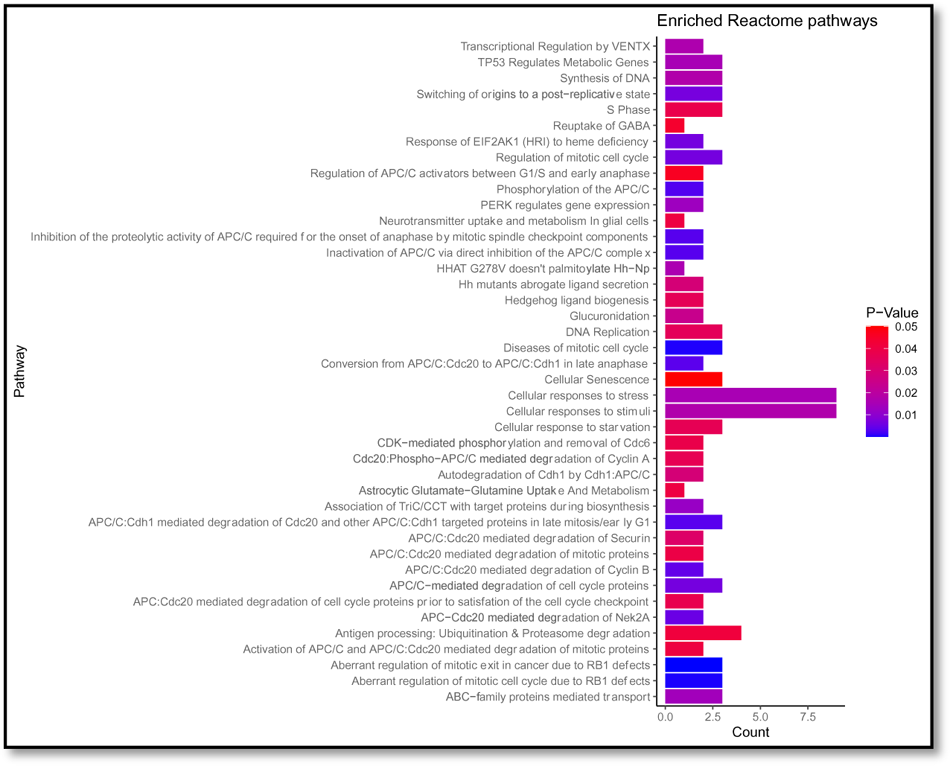
Significantly enriched Reactome pathways of PLMNET mRNAs.

As we see in Fig 5 most PLMNET pathways are related to Anaphase promoting complex/cyclosome (APC/C) and dysregulation of mitotic cell cycle. The APC, a multi-subunit ubiquitin ligase, facilitates mitotic and G1 progression and is now recognized to play a role in maintaining genomic stability. It is also possible that the APC itself plays a crucial role in tumorigenesis through its regulation of mitotic progression. Furthermore, modulating a specific interaction of the APC/C may be therapeutically attractive in specific cancer subtypes, and therapeutic interventions affecting APC/C activity may be beneficial in cancers that are resistant to classical chemotherapy [23].

SKP2, ANAPC2, and ANAPC10 are APC/C substrates with significant differential expression among PRAD tumor and normal samples. These RNAs also have a strong expression correlation with PLMNET hubs.

The APC/C substrate, Skp2, controls the G1 to S transition by eliminating numerous regulatory proteins that inhibit S phase entry. Disruption of signaling pathways controlling the APC/C–Skp2 axis during neuronal differentiation may lead to disruption of homeostasis [24]. Furthermore, proved that Skp2 plays a critical role in the development and progression of prostate cancer. Thus, Skp2 was introduced as a significant therapeutic target in prostate cancer [25]. Ultimately, our analysis illustrates that PLMNET hubs also may represent an attractive therapeutic target in PRAD.

### 3.5. PRAD-specific lncRNAs and overall survival

Kaplan–Meier and log-rank tests were used to explore whether the expression of each lncRNA in PLMNET had prognostic significance for predicting overall survival. Statistical significance was set at p-value <0.05. Finally, 11 lncRNAs, LINC02137, RP11-431N15.2, LINC01671, LINC02806, ENSG00000231196, ENSG00000244252, GATA3-AS1, RP11-379F12.3, LINC01116, EMX2OS and SLC2A9-AS1 were found to be related to OS (Fig 6).

**Fig 6.**
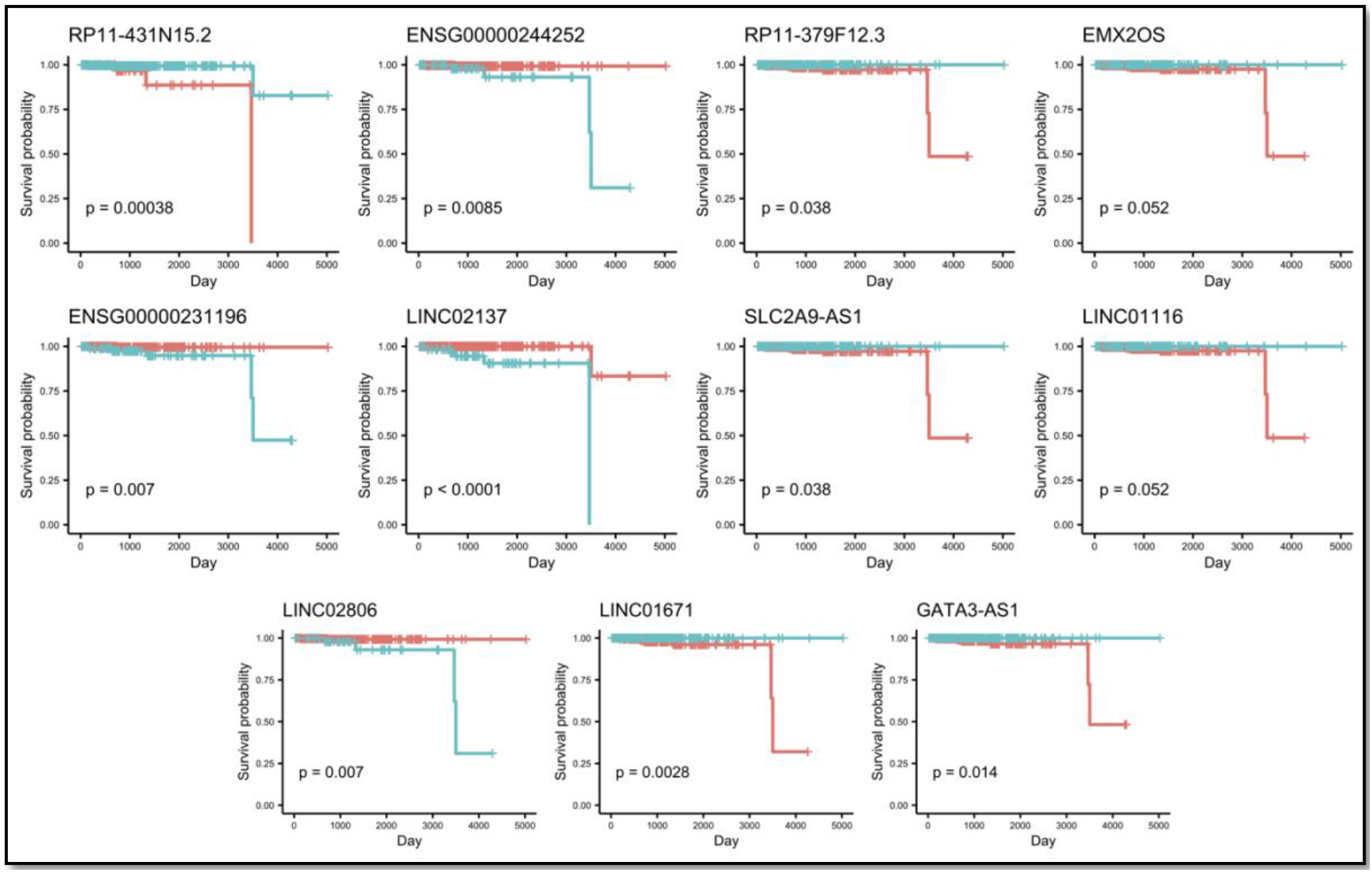
Kaplan–Meier survival curves for PLMNET lncRNAs associated with OS of the PRAD patients. Log-rank method was used to assess the survival differences between the two groups. Horizontal axis is OS time (days), and vertical axis stands for survival function. The turquoise lines represent the group with low-count, and the red lines represent the group with high-count.

### 3.6. Identification of PLMNET lncRNAs associated hallmarks

Although the biology of cancer is extremely complex, the complexity of cancer can be reduced and represented by a few cancer hallmarks. In this part, we focused on the lncRNAs regulations in the context of cancer hallmarks. This leads us toward a better understanding of their role and functions in cancer biology. Recent studies associated some genes with cancer hallmarks. Hallmark genes were regulated by more ncRNAs than others, suggesting that they are more likely to be precisely expressed under the strict regulatory control of ncRNAs [26].

PHANET illustrates the association between PLMNET lncRNAs and cancer hallmarks. It also determined the most significant PRAD hallmark genes which had a strong expression correlation with PLMNET lncRNAs (Supplementary fileFile 6). PHANET consists of 22 lncRNAs, 19 mRNAs, 23 GO-terms, and 8 hallmarks (Fig 7). 26% of PHANET hallmark genes are oncogene or tumor suppressors. The main network is depicted in four different nets for more clarity, Those RNAs that were not connected to any hallmark were omitted from the network. PHANET is a beneficial general overview of most altered RNAs in PRAD tissues and their relation to cancer hallmarks.

**Fig 7.**
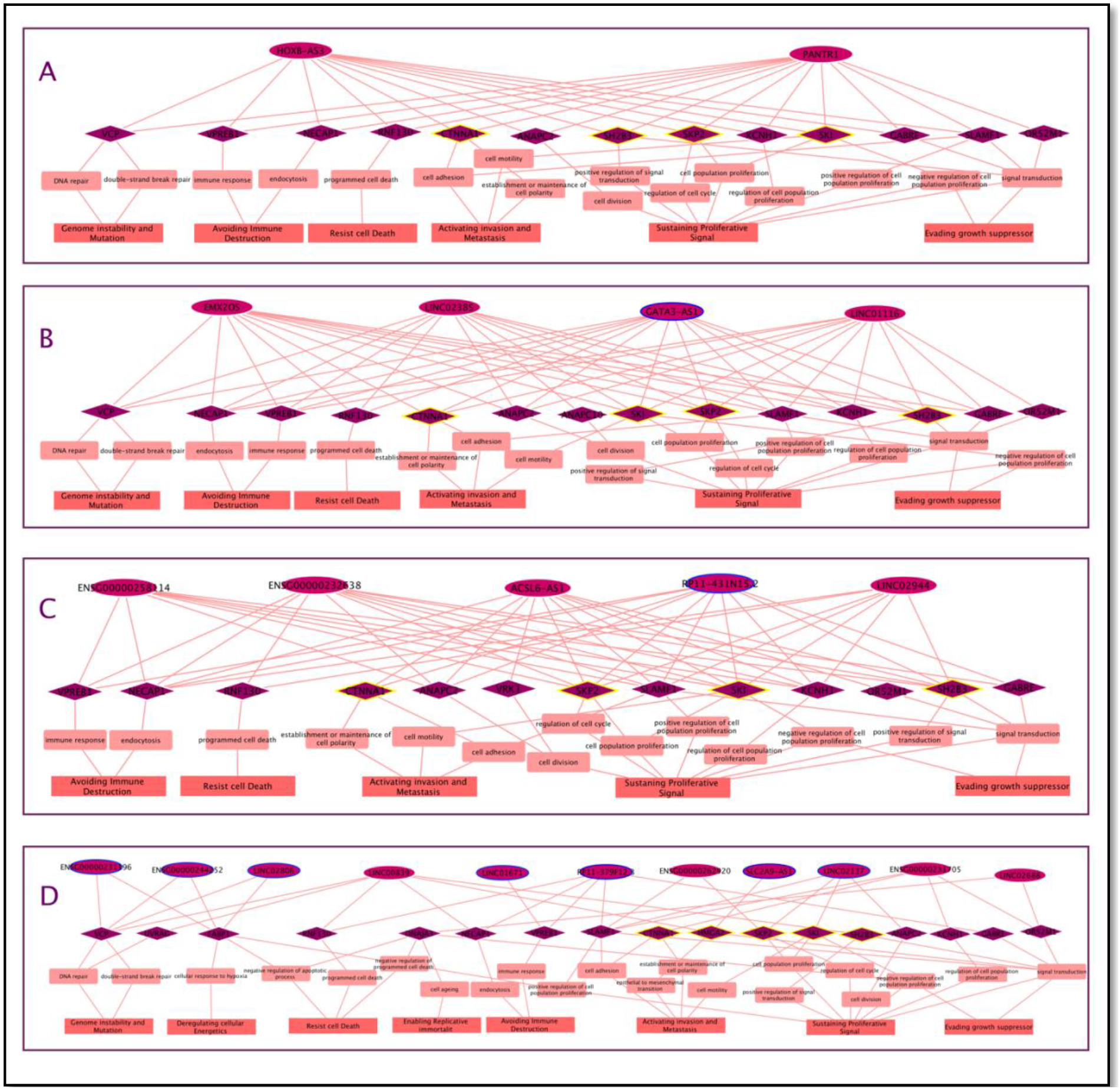
PHANET: PRAD lncRNA-mRNA-GO-Hallmark net. Ellipse: lncRNAs, diamond: mRNAs, salmon rectangles: GO-terms, imperial rectangles: Cancer hallmarks. Yellow borders represent oncogenes or tumor-suppressors and blue borders represent OS-related lncRNAs.

Net A represents the association among the two most high degree lncRNAs of PLMNET, PANTER (degree:32) and HOXB-AS3 (degree:31). The expression correlation among these lncRNAs is 0.979. They also have a high expression correlation with 29 common mRNAs of PLMNET and 12 common mRNAs of PHANET. Among these common mRNAs, SKI and

SKP2 are oncogenes, and SH2B3 and CTNNA1 are tumor suppressors. Most other PHANET mRNAs also have a critical role in cancer. Net B represents the rest of PLMNET hubs (degree 30) and their connection to hallmark genes and cancer hallmarks.

All the hubs have the same connection to 6 cancer hallmarks. Fig 8 depicts common mRNAs (with high expression correlation) among hubs. Fig 7 illustrates PLMNET hubs’ strong regulatory role in PRAD proliferation, invasion, and metastasis, according to the degree of cancer hallmark and their associated GO-terms. They also could interfere with genome instability and mutation, immune destruction, resist cell death and evade growth suppression.

**Fig 8.**
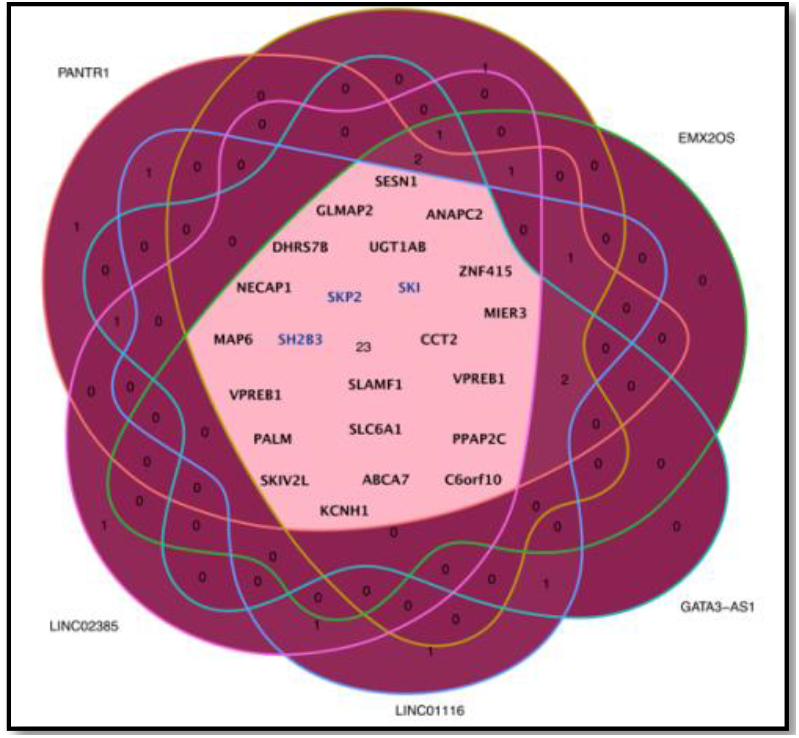
Common mRNAs (with high expression correlation) among PLMNET hubs.

The rest of PLMNET lncRNAs depicts in net C and D. Net D holds lncRNAs with the lowest degrees. In these parts, lncRNAs seem to form distinct functions.

According to all interactions of PHANET, sustaining proliferative signal (via 8 GO-terms), Activating invasion and metastasis (via 4 GO-term), and resist cell death (via 3 GO-term) are the most significant cancer hallmarks of PLMNET.

## 4. Discussion

Transcriptional perturbations are widely mediated by lncRNAs. They are regulators of gene expression and have been shown to play critical roles in cancer. lncRNAs contribute to various cellular processes, such as proliferation, invasion, metastasis, and apoptosis, but yet their molecular mechanisms in PRAD have not been fully studied. The current study investigated the regulation role of lncRNAs in PRAD. The PLMNET was constructed based on RNAs, differentially expressed among tumor and their adjacent normal samples, and had a strong expression correlation with others. This leads to construction of a network with 23 lncRNAs and 52 mRNAs. As depicted in previous parts, all PLMNET lncRNAs could have a key regulatory role in PRAD. Then we focused on the most significant lncRNAs. According to network analysis principles, the top 8% high degree nodes of PLMNET were considered as most significant lncRNAs in PRAD (PANTR1, HOXB-AS3, EMX2OS, GATA3-AS1, LINC01116, LINC02385). Literature review proves that PANTR1, EMX2OS, LINC01116 are key regulatory lncRNAs of prostate adenocarcinoma and HOXB-AS3 and GATA3-AS1 have regulatory roles in other cancer. PANTR1, the highest degree nodes of PLMNET, were highly connected to sustaining proliferative signal and Activating invasion and metastasis hallmarks according to PHANET.

PANTR1 is an oncogenic lncRNA with notable influence on numerous cellular features in different types of cancer. PANTR1 overexpression is recognized in prostate carcinoma and its expression levels increase with the increase in the diameter of tumor. It was demonstrated that PANTR1 may promote cancer cell proliferation in prostate carcinoma by upregulating ROCK1 [27]. Sales et al. also provides first evidence that PANTR1 has a relevant role in human Renal cell carcinoma (RCC) by influencing apoptosis and angiogenesis [28]. PANTR1 can also promote the expression of MDR and stem cell markers in chronic myeloid leukemia cell line K562, and interfere imatinib resistance [29]. PANTR1 promotes Hepatocellular carcinoma (HCC) progression via mediating the miR-587-BCL2A1 axis [30].

HOXB-AS3, the second-high degree node of PLMNET, also had the same connection to cancer hallmarks as PANTR1. HOXB-AS3 is a tumor suppressor downregulated in highly metastatic and primary Colorectal cancer (CRC) tissues. HOXBAS3 inhibits cancer cell proliferation, invasion, and metastasis and suppresses tumor growth. The loss of HOXB-AS3 is a significant oncogenic event in CRC [31]. Furthermore, HOXB-AS3 is overexpressed in Endometrial carcinoma (EC) tissues and cell lines. Interference with HOXB-AS3 expression can inhibit endometrial carcinoma EC cell proliferation and promote apoptosis [32]. Inhibition of HOXB-AS3 expression in Hepatocellular carcinoma (HCC) was confirmed to promote proliferation and inhibit apoptosis. This mechanism was associated with the regulation role of HOXB-AS3 in p53 expression by binding to DNMT1[33].

EMX2OS was identified as the enhancer RNA (a subclass of lncRNAs transcribed from enhancer regions that play an important role in the transcriptional regulation of genes) in kidney renal clear cell carcinoma. EMX2OS downregulation was significantly associated with higher histological grade, advanced stage, and poorer prognosis [34]. EMX2OS also were downregulated in prostate cancer tissues and cells and may be regarded as a potential molecular biomarker for diagnosis and prognosis. EMX2OS negatively regulated cell proliferation, migration, and invasion [35]. This evidence confirms our findings about the role of this lncRNA in prostate cancer.

LINC01116 (another hub of PHANET) expression dysregulation is correlated with a variety of cancers, including gastric cancer, lung cancer, colorectal cancer, glioma, and osteosarcoma. LINC01116 plays a crucial role in facilitating cell proliferation, invasion, migration, and apoptosis and is suggested as a novel biomarker for prognosis and therapy in malignant tumors [36]. LINC01116 also is involved in the development of osteosarcoma by binding to EZH2 to regulate expressions of PTEN and p53 [37]. Furthermore, Functional assays indicated that inhibition of LINC01116 could suppress cell proliferation, migration, invasion, and Epithelial-mesenchymal Transition progress in prostate cancer cells. Yu et al. suggested that LINC01116 acted as an oncogene in prostate cancer and accelerated prostate cancer cell growth through regulating miR-744-5p/UBE2L3 axis [38]. According to our enriched Reactome pathways, PLMNET was associated with “TP53 Regulates Metabolic Genes” pathway via SESN1(P-value = 0.01). SESN1 also had a high expression correlation with 13 lncRNAs of PLMNET. Six hubs of PLMNET are among those 13 lncRNAs. Further functional experiments will be required to validate the effect of PLMNET hubs on SESN1 and P53 and vice versa in PRAD. GATA3-AS1 promotes cell proliferation and metastasis of HCC by suppression of PTEN, CDKN1A, and TP53. GATA3-AS1 overexpression was significantly correlated with larger tumor size, advanced TNM stage, more lymph node metastasis, and OS [39]. It also modulates tumorigenesis in pancreatic cancer, which may be associated with the Wnt/β-catenin signaling pathway [40]. GATA3-AS1 contributed to Triple negative breast cancer (TNBC) progression and immune evasion through stabilizing PD-L1 protein and degrading GATA3 protein, offering a new target for the treatment of TNBC [41].

GATA3-AS1 was established as an independent predictor of response to neoadjuvant chemotherapy in Breast cancer and proposed as a potential predictive biomarker of nonresponse to neoadjuvant chemotherapy [42]. The result of Kaplan–Meier survival analysis in previous parts also connects the GATA3-AS1 high expression to low OS.

Among all hubs of PLMNET, yet there is no evidence of LINC02385 participation in cancer, but according to its high degree and BC, we expect to explore its contribution in early future. Ultimately, PLMNET hubs seem to contribute to PRAD proliferation, invasion, metastasis, and apoptosis. PLMNET hubs also may be regarded as a potential molecular biomarker for diagnosis and prognosis of PRAD and develop therapeutic targets for PRAD patients.

Despite the significant key regulatory role of PLMNET hubs in PRAD or other cancers, yet we expect other PLMNET lncRNAs to have a key regulatory role in PRAD because of their differential expression among tumor and normal tissues, correlation with OS, and hallmark genes.

## 5. Conclusion

Evidence accumulated over the past decade suggests that dysregulation of lncRNA is ubiquitous in all human tumors. lncRNAs contribute to various cellular processes, such as proliferation, invasion, metastasis, and apoptosis. The current study comprehensively investigated the regulatory role of lncRNAs in PRAD. The PRAD lncRNA-mRNA network was constructed based on RNAs that were differentially expressed among tumor and normal samples and had a strong expression correlation with others. Our results proved that hubs of network have an association with overall survival and their regulatory roles have been confirmed in diverse cancer. Therefore, these hubs may be regarded as a potential molecular biomarker for diagnosis and prognosis of PRAD and develop therapeutic targets for PRAD patients.

## Abbreviations

lncRNAs: Long non-coding RNAs
PRAD: Prostate adenocarcinoma
PLMNET: lncRNA-mRNA network
APC/C: Anaphase promoting complex/cyclosome
TCGA: The Cancer Genome Atlas
RSEM: RNA-Seq by Expectation Maximization
TANRIC: The Atlas of Noncoding RNAs in Cancer
RPKM: Reads per kilo base per million mapped reads
BC: betweenness centrality
OS: Overall Survival
PHANET: PRAD hallmark associated RNA network
GO: Gene Ontology
RCC: Renal cell carcinoma
HCC: Hepatocellular carcinoma
CRC: Colorectal cancer
EC: Endometrial carcinoma
TNBC: triple negative breast cancer

## Availability of data and materials

The mRNA expression profiles were downloaded from TCGA (https://portal.gdc.cancer.gov) and lncRNAs expressions were directly obtained from ‘The Atlas of Noncoding RNAs in Cancer’(TANRIC, https://ibl.mdanderson.org/tanric/_design/basic/download.html). The rest of the information is included within the article and its additional files.

## Authors Contribution

HR conceived the research. HR and AK designed the research. HR implemented the research and wrote the manuscript. All authors read and approved the final manuscript.

