## Supplementary material for "lncRNA-mRNA network analysis reveals novel biomarkers in prostate adenocarcinoma": List of supplementary files.docx

S1: The most significant differentially expressed mRNAs.

S2: The most significant differentially expressed lncRNAs.

S3: List of PLMNET edges with their correlations and P-values.

S4: PLMNET node list with degrees and BCs.

S5: 41 Reactome pathways associated with PLMNET mRNAs.

S6: Association between lncRNAs and Cancer hallmarks.
